# conMItion: an R package adjusting confounding factors for associations in multi-omics

**DOI:** 10.64898/2026.05.07.723535

**Authors:** Gaojianyong Wang, Frank Liu, Ze Chen, Teresa Davoli

## Abstract

**Summary:** Association measurements, such as mutual information (MI), are fundamental in the analysis of cancer multi-omics data for identifying cancer-related genes, gene signatures, and gene regulatory networks, thereby shedding light on tumor development, progression, and treatment. Confounding factors, including tumor purity and mutation burden, can bias association measurements in MI, potentially leading to the misclassification of passenger events as drivers. Conditional mutual information (CMI) provides a robust framework for assessing both linear and non-linear associations while effectively accounting for different confounding factors. An R package called `conMItion` is introduced to estimate CMI and its statistical significance for multi-omics data, with flexibility to adjust for one or two confounding factors. We demonstrated the utilization of `conMItion` through two use cases. First, we identified co-occurring somatic alterations in bladder cancer genomic data. Second, we applied `conMItion` to a single-cell RNA sequencing dataset of lung cancer patients and identified positively or negatively associated cell types within the lung cancer tumor microenvironment.

**Availability and Implementation:** The `conMItion` package is freely available on CRAN at https://CRAN.R-project.org/package=conMItion. The two use cases described in the paper can be accessed at https://github.com/GJYWang/conMItion. A supplementary document is available online.

## Introduction

Association measurement is one of the most fundamental tools for multi-omics analyses in cancer research, shedding light on the complex interactions driving tumor development and progression. For example, copy-number-dependent oncogenes and tumor suppressor genes have been identified by examining the associations between somatic copy number alterations (SCNAs) and gene expression (1). Co-expressed gene signatures have also led to the identification of biomarkers linked to cancer hallmarks (2). In addition, association measurements can reveal gene regulatory networks (3), providing insights into new drug targets.

Mutual information (MI) is a robust association measurement method for quantifying both linear and non-linear dependencies in the intricate interactions in cancer (4). However, confounding factors such as tumor purity (5) and mutation burden (6) can significantly bias association analyses (7). If these confounding factors are not properly addressed, the identification of tumor drivers through association analyses may mistakenly identify passenger events associated with tumor purity or mutation burden as tumor drivers.

Therefore, addressing these confounding factors is essential for improving the accuracy and reliability of biological inferences obtained from association measurements. Conditional mutual information (CMI) serves as an ideal framework to address different confounding factors (8), as it evaluates both linear and non-linear associations while theoretically accounting for an unlimited number of confounding factors. Therefore, we have developed an R package called `conMItion` to estimate CMI for biological data. In this paper, we present two illustrative use cases of `conMItion`, demonstrating its practical application. The first case is the identification of co-occurring mutations and SCNAs while controlling for tumor purity and mutational burden as confounding variables. The second case is the evaluation of the association between the abundance of different cell types within the tumor microenvironment (TME), while controlling for tumor purity as a confounding variable.

## Methods

### 2.1 Information theory basics

Entropy quantifies the amount of information needed to describe a random variable. For a discrete random variable *X* with probability mass function *p*(*x*) over possible outcomes 𝒳, the entropy is defined as:

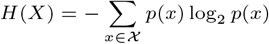

The base-2 logarithm ensures that entropy is measured in bits. The joint entropy quantifies the amount of information required to describe two or more random variables. For two discrete random variables *X* and *Y* with joint probability mass function *p*(*x, y*) over all possible outcomes 𝒳 *×* 𝒴, the joint entropy is defined as:

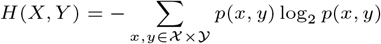

For three variables *X, Y*, and *Z*, with joint probability mass function *p*(*x, y, z*) over all possible outcomes 𝒳 *×* 𝒴 *× Ƶ*, the joint entropy is defined as:

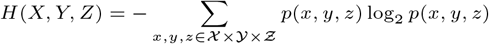

Mutual information (MI), *I*(*X*; *Y*), quantifies the information shared between two random variables *X* and *Y*. It represents the reduction in uncertainty of *X* given *Y* and is expressed as:

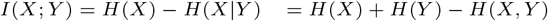

Conditional mutual information (CMI), *I*(*X*; *Y* |*Z*), quantifies the information shared between *X* and *Y* given a third condition variable *Z* (Figure 1A). It is defined in terms of entropy as:

**Fig. 1.**
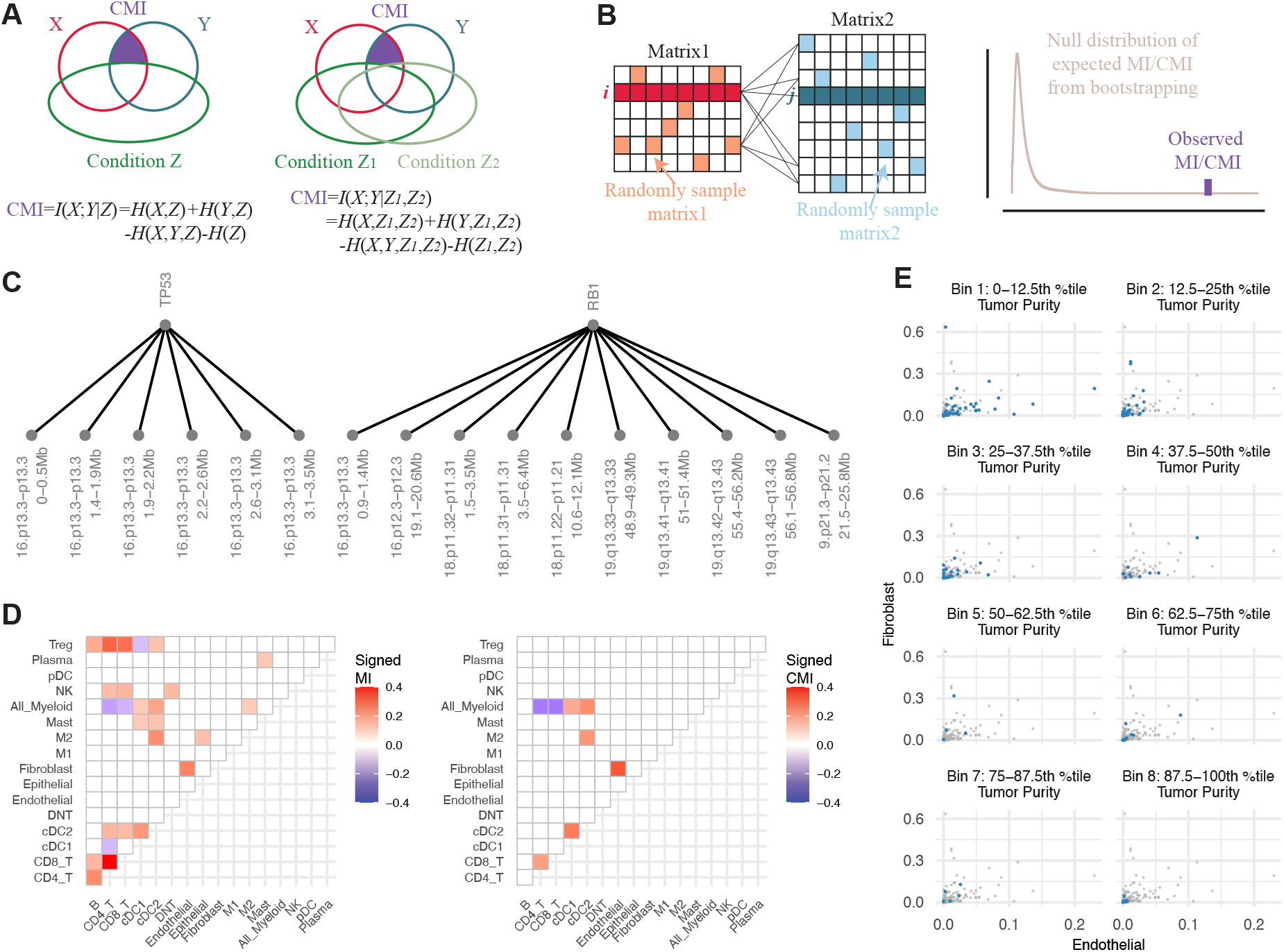
The mathematical fundamentals, workflow, and results of the conMItion package. (A) Venn diagram illustrating CMI between variables *X* and *Y* given the condition variable *Z* (left panel) and given condition variables *Z*_1_ and *Z*_2_ (right panel). The red and blue circles respectively represent the entropy *H*(*X*) and *H*(*Y*). The intersection of the red and blue circles represent MI *I*(*X*; *Y*). The shaded area (purple) represents CMI *I*(*X*; *Y* |*Z*) (left panel) and *I*(*X*; *Y* |*Z*_1_, *Z*_2_) (right panel). (B) Illustration of `conMItion` estimating statistical significance through bootstrapping. (C) SCNAs identified to be associated with mutations of TP53 and RB1 with *P* values less than 1 *×* 10^−3^ (before adjustment for multiple comparisons). Genes are illustrated at the top, and genomic regions are illustrated at the bottom. (D) Statistically significant (*P <* 5 *×* 10^−2^ before adjustment for multiple comparisons) association between cell fractions within lung adenocarcinoma (MI: left panel, CMI: right panel) with malignant cell fraction as the confounding factor. The sign was given by Spearman correlation. (E) Scatter plots of endothelial and fibroblast cell fractions within lung adenocarcinoma with malignant cell fraction (tumor purity) as the confounding factor. The tumor purity is split into 8 bins.

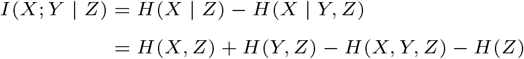

CMI evaluates the dependency between *X* and *Y* while excluding the influence of *Z*. It can be extended to account for multiple conditions. For condition variables *Z*_1_ and *Z*_2_, the CMI, *I*(*X*; *Y* |*Z*_1_, *Z*_2_) (Figure 1A), is defined in terms of entropy as:

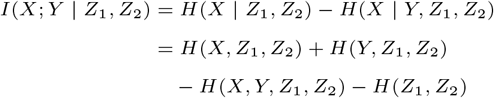

### 2.2 CMI estimation in multi-omics

Marginal and joint probabilities must be estimated before calculating the CMI in multi-omics data. For a variable *X* with observed values *x*_1_, …, *x*_*N*_, these values are normalized between 0 and 1:

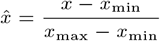

The range between 0 and 1 is divided into *B* bins of the same size, represented by 𝒳_1_, …, 𝒳_*B*_. B-spline functions (4) are used to estimate the probability mass function of *p*(𝒳) (Supplementary Document). A knot vector *t*_*i*_ is defined by a number of bins *i* = 1, …, *B* and a spline order of integer *s*, 0 *< s < B*:

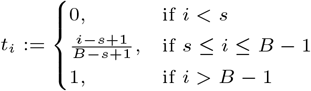

B-spline basis functions 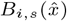 are recursively constructed (Supplementary Document) as follows:

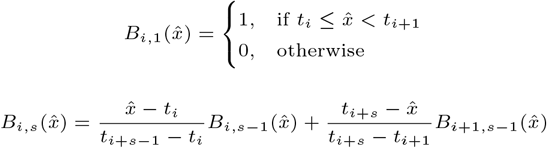

The probability mass function is then estimated as:

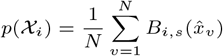

Joint probabilities for two and three variables are respectively given as:

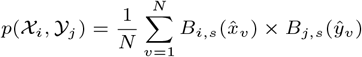

and

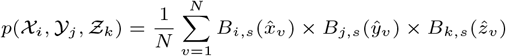

CMI can then be estimated using information theory principles as outlined in Section 2.1. This method also applies to two or more condition variables. The obtained MI or CMI is normalized by max(*I*(*X*; *X*), *I*(*Y*; *Y*)) to have a maximum value of 1 and a minimum value of 0.

### 2.3 conMItion package

`conMItion` has been developed for association measurement in multi-omics data and for assessing the statistical significance of the obtained associations. The package accommodates different data formats and facilitates the distribution of large computational tasks across multiple nodes. The functions are divided into two main categories. The first set of functions estimates MI and CMI for different data formats, such as between a matrix and a vector or between two matrices. The MI and CMI values are normalized to range from 0 to 1, facilitating comparisons across different datasets. The second set of functions generates distributions of MI/CMI association measures through bootstrapping (Figure 1B). By randomly sampling input matrices, the package computes a null distribution of MI/CMI association measures. Statistical significance can be obtained by comparing observed associations against the context of the null distribution.

## Results

### 3.1 Co-occurring mutations and SCNAs

`conMItion` was applied to Urothelial Bladder Carcinoma (BLCA) samples from TCGA to identify co-occurring missense mutations and SCNAs. We collected data on the frequency of missense mutations for 20,724 genes, and obtained the average SCNA levels for 1,368 genomic segments that together comprise the whole genome (Supplementary Methods). Across 393 patient samples, `conMItion` estimated the CMI between the mutational status of 20,724 genes and the SCNAs of 1,368 genomic segments while conditioning on tumor purity and mutation burden, using a bin number of 8 and a spline order of 2 (Supplementary Document). There were 374 mutation-SCNA pairs whose CMI had *P <* 10^−3^ (before adjustment for multiple comparisons). We found that TP53 mutations co-occurred with with 16p13.3 SCNAs (Figure 1C). 16p13.3 contains gene E4F1, a transcription factor known to interact with TP53 (9). In this region, 29.0% BLCA samples showed deletions smaller than −0.2 (log2 ratio). Additionally, 21.4% of BLCA samples acquired both TP53 mutations and E4F1 deletions.

Furthermore, `conMItion` also evaluated the CMI between gene mutations and segment SCNAs conditioned only on tumor purity (termed CMI_*p*_ here to differentiate it from CMI conditioned on both tumor purity and mutation burden, or CMI_*p,m*_, as calculated above). The association between GINS3 missense mutation and SCNA at 2q34, containing ERBB4, reached statistical significance when overall mutation burden was not controlled (MI = 0.089, *P* = 5.4 *×* 10^−4^ pre-adjustment; CMI_*p*_ = 0.065, *P* = 1.8 *×* 10^−2^ pre-adjustment). However, CMI_*p,m*_ for this mutation-SCNA pair was 0.028 (*P* = 0.20 pre-adjustment), suggesting the magnitude of the association was inflated by mutational burden as a confounding variable. Similarly, the association between EGFR3 missense mutations and SCNAs at 9q21.33 (MI = 0.087, *P* = 6.2 *×* 10^−4^ pre-adjustment) underwent a decrease in magnitude after conditioning on tumor purity (*CMI*_*p*_ = 0.046, *P* = 0.055 pre-adjustment), suggesting that the observed MI association for this mutation-SCNA pair could be inflated by tumor purity. These examples demonstrate how `conMItion` can distinguish between genuine co-occurring genomic alterations of interest and spurious associations driven by confounding variables, improving the accuracy of driver gene identification.

### 3.2 Cell Type Associations in the TME

Twelve single-cell RNA sequencing (scRNA-seq) datasets of lung cancer were collected and processed. Cell type annotation was performed through the detection of canonical marker genes, and the fractional abundance of every cell type within the TME was subsequently computed across all samples (Supplementary Methods).

Signed MI and CMI (conditioned on tumor purity) were calculated between all cell type fractions using bin of 8 and spline order of 2. The association sign was set as the sign of the corresponding Spearman correlation (Figure 1D). Co-occurring cell type pairs included CD4^+^ T cells with CD8^+^ T cells (CMI = 0.22, *P* = 3.4 *×* 10^−3^ pre-adjustment), and endothelial cells (ECs) with fibroblasts (FIBs) (CMI = 0.32, *P* = 3.9 *×* 10^−6^ pre-adjustment, Figure 1E). Additionally, associations between Treg fraction and various lymphocytes (B cells, CD4^+^ T cells, and CD8^+^ T cells) that were found to be statistically significant by MI (0.17, 0.31, 0.29 respectively; *P <* 10^−3^, 10^−6^, 10^−6^ respectively) became insignificant after conditioning on tumor purity (CMI = 0.11, 0.14, 0.15 respectively; *P* = 0.56, 0.20, 0.13 respectively), suggesting the magnitude of these associations may be confounded by tumor purity. This finding makes biological sense, as Tregs are responsible for supression of immune responses, rather than direct participation in antitumor cytotoxicity. Their association with other lymphocytes appears to be a side effect of overall tumor purity, rather than specific cellular interactions with B cells or conventional T cells. This use case illustrates how `conMItion`’s ability to condition on confounding factors like tumor purity can both highlight previously obscured associations (endothelial-fibroblast) and filter out spurious correlations (Treg-lymphocyte), providing a more nuanced understanding of cell type co-localization patterns in the TME.

## Discussion and Conclusion

This paper presents the R package `conMItion`, which uses CMI to control for the effects of confounding factors in association measurements in multi-omics analyses. Through two use cases, we demonstrated that `conMItion` offers additional insights into the complex genomic alteration patterns and cell type co-localization patterns in cancer. In the first use case, we found that TP53 missense mutations and somatic deletions of E4F1 tend to co-occur in BLCA (after adjusting for both tumor purity and mutation burden), which is consistent with the known complementary roles played by the two transcription factors in regulating cell cycle progression (9).

In the second use case, we found that the positive association between FIBs and ECs (10) in the TME becomes more apparent after accounting for tumor purity. This finding is biologically meaningful, as cancer-associated FIBs are known to promote angiogenesis and support EC function through the secretion of pro-angiogenic factors such as VEGF, FGF2, and chemokines (11). Moreover, ECs can undergo endothelial-to-mesenchymal transition (EndMT) to become a source of cancer-associated FIBs in the TME, providing a direct mechanistic link between these two cell types (12). The stronger CMI compared to MI indicates that this stromal interaction occurs consistently across varying levels of tumor purity, suggesting it represents a fundamental feature of tumor-stroma crosstalk rather than an artifact of tumor cell content.

Importantly, our analysis also revealed that associations between Tregs and other lymphocyte populations (B cells, CD4^+^ cells, CD8^+^ T cells) that appeared significant by MI became non-significant when calculating CMI, indicating that these associations were largely driven by tumor purity rather than specific cellular interactions. This observation is biologically plausible, as Tregs are a specialized immunosuppressive subset of CD4^+^ T cells characterized by FOXP3 expression, functioning to suppress antitumor immunity rather than directly collaborating with conventional effector lymphocytes. While Tregs normally comprise only 4% of CD4^+^ cells in peripheral blood, they can constitute 20–30% of CD4^+^ cells in the TME. Their accumulation in tumors occurs through preferential chemotaxis (e.g., via CCR4-CCL22 gradients) and reflects their role in creating an immunosuppressive microenvironment enabling tumor immune evasion (13). Thus, the apparent associations between Tregs and other lymphocytes detected by MI likely reflect co-enrichment patterns related to overall immune infiltration (inversely correlated with tumor purity) rather than functional co-localization or direct cellular interactions.

Together, these examples demonstrate how `conMItion` can distinguish between genuine biological associations of interest versus spurious correlations driven by confounding variables, and underscore the importance of proper confounder adjustment in multi-omics association analyses.

## Author contributions statement

G.W. wrote and maintained the package. G.W., F.L., Z.C., and T.D. analysed the results, G.W., F.L., Z.C., and T.D. wrote the manuscript.

## Acknowledgments

The author extends special gratitude to Marisa Korody, Romualdas Smičius, and Martin Vingron for their invaluable discussions and support.

