## Supplementary Material for "conMItion: an R package adjusting confounding factors for associations in multi-omics"

This supplementary document provides supplementary methods and additional analysis on the use of mutual information (MI) and conditional mutual information (CMI) for measuring associations in multi-omics data. This document serves to supplement the main paper by offering additional insights and supporting evidence for the arguments and findings presented therein.

The document begins with supplementary methods for Section 3.1 and Section 3.2 from the main manuscript, and the full list of top co-occurring mutation-SCNA pairs in bladder cancer.

Next, the document summarizes an extended introduction to B-spline functions by providing illustrated examples of how knot vectors and B-spline functions are generated, which are core steps in the protocol for non-parametric estimation of MI and CMI.

Furthermore, using data from Section 3.1, we investigate the impact of tuning bin number and spline order hyperparameters on the magnitude of the estimated MI/CMI and associated $p$-values. Our results align with previous analyses, which have established MI estimation based on B-spline functions as a robust association measurement tool [1].

Lastly, using data from Section 3.2, we present a comparative analysis of MI and CMI with correlation and linear regression. We illustrate the advantages and limitations of each approach for discovering complex associations in biological data.

Together, these sections provide a comprehensive exploration of MI and CMI estimation using B-splines for biological data, offering readers both concrete evidence and practical guidance for utilizing the developed R package `conMItion`.

### 1 Supplementary Methods

#### 1.1 Bladder Cancer Genomic Data

`conMItion` was applied to Urothelial Bladder Carcinoma (BLCA) samples from TCGA to identify co-occurring missense mutations and SCNAs. The mutation data for 20,724 genes were obtained from the Multi-Center Mutation Calling in Multiple Cancers (MC3, [2]), while SCNA data generated by the Affymetrix SNP 6.0 platform were obtained via TCGAbiolinks [3]. The total number of missense mutations for all genes was used to represent mutation burden. Harmonized expression data were downloaded via TCGAbiolinks [3] to obtain transcript information. Roughly 54,000 transcripts were downloaded (detailed transcript information can be accessed at `https://github.com/GJYWang/conMItion`). To reduce computational time, 40 adjacent transcripts within the same chromosome arm, or remaining transcripts at the end of a chromosome arm, were merged into segments, resulting in 1,368 fragments. The SCNA level of a segment was calculated as the average SCNA of its transcripts. Tumor purity data for BLCA samples were obtained from a recent publication [4]. A total of 393 BLCA samples with mutation, SCNA, and tumor purity information were analyzed. See Table 1 below for the full list of discovered mutation-SCNA associations.

#### 1.2 Single-cell RNA Sequencing Dataset of Lung Cancer

Six single-cell RNA sequencing (scRNA-seq) datasets of lung cancer, sequenced using the 10X Genomics platform, were sourced from the Curated Cancer Cell Atlas [5]. Additional datasets were obtained from repositories including the Gene Expression Omnibus (GSE148071), the Sequence Read Archive (PRJNA634159, PRJCA001731, PRJNA622993, PRJNA1055415), and the ArrayExpress database (E-MTAB-6149, E-MTAB-6653). Seurat [6] was used to process all scRNA-seq data, applying dataset-specific quality control thresholds for metrics such as the number of detected genes and the total number of cells. Integration across datasets was performed using Harmony. Cell type annotation was performed through the detection of canonical marker genes (see Supplementary Methods for details). The number of cells for each cell type was quantified for all samples, and cell fraction vectors were computed by normalizing cell counts by total cell quantity per sample. Tumor purity was defined as the malignant cell fraction per sample.

#### 1.3 Cell Type Annotation

Cell type annotation was performed as follows: NK cells were identified based on the expression of KLRC1, KLRD1, and NKG7; $CD8^+$ T cells were annotated using CD3D, CD3E, CD3G, CD8A, GNLY, and GZMA; $CD4^+$ T cells were annotated using CD4; Treg cells were annotated with FOXP3,CTLA4; B cells were defined by MS4A1, CD79A, and CD79B; plasma cells were defined with CD38,SDC1,JCHAIN; fibroblasts were defined by COL1A1, COL1A2, and DCN; endothelial cells were identified by CLDN5 and VWF; myeloid cells except DC were defined by CD14, LYZ, CD68; M1 macrophages were defined by CD64, CD80, CXCL9, CXCL10, CXCL11, CD86, IL1A, IL6, CD40; M2 macrophages were defined by ARG1, ARG2, IL10, CD163, CCL4, CCL13, CCL17, CCL20, MRC1, MSR1; pDC were annotated by IL3RA, LILRA4, CLEC4C; cDC1 were defined with XCR1, CLEC9A; cDC2 were defined with FCER1A,CD1C; malignant cells were identified by NAPSA, TTF1, KRT5, DSG1, TP63; epithelial cells were annotated by using CAPS, SNTN; and mast cells were defined by GATA2, TPSAB1, TPSB2.

### 2 Top co-occurring mutation-SCNA pairs

Table 1: Top co-occuring mutation-SCNA pairs in BLCA with P<0.001

| Mutation | SCNA | $CMI_{p,m}$ | P Value |
|---|---|---|---|
| RPS6 | 1:76.7-78.7Mb | 0.155 | 0.00003 |
| MTA1 | 12:10.6-11.5Mb | 0.155 | 0.00003 |
| DONSON | 12:10.6-11.5Mb | 0.153 | 0.00004 |
| C1orf127 | 12:10.6-11.5Mb | 0.150 | 0.00005 |
| HNRNPA2B1 | 1:121.6-121.6Mb | 0.147 | 0.00005 |
| GPR162 | 1:121.6-121.6Mb | 0.145 | 0.00007 |
| ABL1 | 12:10.6-11.5Mb | 0.144 | 0.00007 |
| HCRTR2 | 7:112.3-116.3Mb | 0.141 | 0.00008 |
| MSI2 | 1:53.9-55.2Mb | 0.138 | 0.00010 |
| PDE5A | 1:121.6-121.6Mb | 0.137 | 0.00011 |
| CEP250 | 1:119.9-121.6Mb | 0.137 | 0.00011 |
| PRPF4B | 1:189.9-193.8Mb | 0.136 | 0.00012 |
| SPEM2 | 1:206.1-207.3Mb | 0.136 | 0.00012 |
| VWCE | 13:48.3-50Mb | 0.136 | 0.00012 |
| MED12 | 4:81.3-83.4Mb | 0.136 | 0.00012 |
| MMP2 | 17:38.4-39.1Mb | 0.135 | 0.00012 |
| SLC25A40 | 12:10.6-11.5Mb | 0.135 | 0.00013 |
| GSTM2 | 1:40.7-42.6Mb | 0.134 | 0.00013 |
| VIL1 | 1:76.7-78.7Mb | 0.134 | 0.00014 |
| RHBG | 6:89.3-93.1Mb | 0.133 | 0.00015 |
| SORBS3 | 12:10.6-11.5Mb | 0.132 | 0.00016 |
| LILRB1 | 11:54.6-55.7Mb | 0.132 | 0.00016 |
| NFATC1 | 1:121.6-121.6Mb | 0.129 | 0.00020 |
| SLCO1A2 | 5:114.5-116.6Mb | 0.128 | 0.00020 |
| LRRC37A3 | 12:64.3-65.8Mb | 0.128 | 0.00021 |
| PUS3 | 1:121.6-121.6Mb | 0.128 | 0.00021 |
| FAM161A | 1:121.6-121.6Mb | 0.128 | 0.00022 |
| CEP250 | 1:121.6-121.6Mb | 0.127 | 0.00022 |
| AZIN1 | 1:76.7-78.7Mb | 0.127 | 0.00022 |
| MSH4 | 12:55.4-56Mb | 0.127 | 0.00023 |
| TPCN1 | 1:121.6-121.6Mb | 0.127 | 0.00023 |
| LIMA1 | 1:121.6-121.6Mb | 0.127 | 0.00023 |
| RFWD2 | 12:55.4-56Mb | 0.126 | 0.00024 |
| MFRP | 1:76.7-78.7Mb | 0.126 | 0.00024 |
| PADI2 | 1:76.7-78.7Mb | 0.126 | 0.00024 |
| MMP2 | 17:49.8-50.5Mb | 0.125 | 0.00025 |
| CDH6 | 17:31.8-32.5Mb | 0.125 | 0.00026 |
| DLEC1 | 1:121.6-121.6Mb | 0.125 | 0.00026 |
| ITGAV | 1:40.7-42.6Mb | 0.125 | 0.00027 |
| SLCO1A2 | 5:100.7-107.2Mb | 0.124 | 0.00027 |
| ICE1 | 22:22.6-22.8Mb | 0.123 | 0.00029 |
| PAN2 | 12:25.1-27Mb | 0.123 | 0.00029 |
| CNTN4 | 1:119.9-121.6Mb | 0.123 | 0.00029 |
| COMP | 6:105.1-107.8Mb | 0.123 | 0.00029 |
| DNAH2 | 1:121.6-121.6Mb | 0.123 | 0.00030 |
| ETAA1 | 2:191-195.6Mb | 0.123 | 0.00030 |
| CFLAR | 1:119.9-121.6Mb | 0.123 | 0.00031 |
| ANGPT1 | 11:59.4-60.4Mb | 0.122 | 0.00031 |
| RPS6 | 1:73.3-76.6Mb | 0.122 | 0.00031 |
| LTBP1 | 1:40.7-42.6Mb | 0.122 | 0.00031 |
| ABCC11 | 5:114.5-116.6Mb | 0.122 | 0.00032 |
| MITF | 17:37-38.3Mb | 0.122 | 0.00032 |
| MDC1 | 1:121.6-121.6Mb | 0.122 | 0.00033 |
| ZFYVE9 | 1:40.7-42.6Mb | 0.121 | 0.00033 |
| TRIM58 | 12:10.6-11.5Mb | 0.121 | 0.00033 |

| | | | |
|---|---|---|---|
| MAOA | 17:37-38.3Mb | 0.121 | 0.00033 |
| ZNF341 | 11:55.7-56.2Mb | 0.121 | 0.00033 |
| PADI1 | 5:114.5-116.6Mb | 0.121 | 0.00033 |
| ELAVL4 | 11:77.1-78.6Mb | 0.121 | 0.00034 |
| TTC21B | 1:76.7-78.7Mb | 0.121 | 0.00035 |
| ZKSCAN1 | 11:55.7-56.2Mb | 0.120 | 0.00035 |
| NPTX1 | 7:112.3-116.3Mb | 0.120 | 0.00036 |
| ZFYVE9 | 1:39.7-40.7Mb | 0.120 | 0.00037 |
| TMEM184A | 12:10.6-11.5Mb | 0.120 | 0.00037 |
| OR8B12 | 12:41.7-43.8Mb | 0.120 | 0.00037 |
| SLC4A1 | 5:114.5-116.6Mb | 0.120 | 0.00037 |
| TMC4 | 12:10.6-11.5Mb | 0.120 | 0.00038 |
| PARP10 | 11:55.7-56.2Mb | 0.119 | 0.00039 |
| UGT1A5 | 1:76.7-78.7Mb | 0.119 | 0.00039 |
| STAG3 | 1:53.9-55.2Mb | 0.119 | 0.00039 |
| LTN1 | 16:35.6-35.9Mb | 0.119 | 0.00039 |
| ARMC2 | 12:10.6-11.5Mb | 0.119 | 0.00040 |
| TRPA1 | 19:27.6-29.5Mb | 0.119 | 0.00040 |
| IGSF9B | 22:22.6-22.8Mb | 0.119 | 0.00040 |
| FRMD6 | 17:48.3-48.9Mb | 0.119 | 0.00040 |
| GMPS | 17:31.8-32.5Mb | 0.119 | 0.00040 |
| ITGA2B | 12:10.6-11.5Mb | 0.119 | 0.00040 |
| RABEP2 | 1:121.6-121.6Mb | 0.119 | 0.00040 |
| MARVELD3 | 17:31.8-32.5Mb | 0.119 | 0.00040 |
| RRP12 | 1:38-39.7Mb | 0.118 | 0.00041 |
| DOCK9 | 19:9.7-10.3Mb | 0.118 | 0.00041 |
| CREB3 | 11:55.7-56.2Mb | 0.118 | 0.00042 |
| LMOD2 | 1:76.7-78.7Mb | 0.118 | 0.00042 |
| ZNF615 | 1:119.9-121.6Mb | 0.118 | 0.00042 |
| MMP2 | 17:37-38.3Mb | 0.118 | 0.00043 |
| SMC2 | 22:22.8-23.4Mb | 0.118 | 0.00043 |
| ESRP1 | 2:191-195.6Mb | 0.118 | 0.00043 |
| SYT3 | 13:48.3-50Mb | 0.118 | 0.00043 |
| RIPK4 | 4:10.5-15.1Mb | 0.117 | 0.00044 |
| ALDOA | 1:121.6-121.6Mb | 0.117 | 0.00044 |
| MAGEC3 | 12:65.8-68.2Mb | 0.117 | 0.00044 |
| ZNF626 | 1:189.9-193.8Mb | 0.117 | 0.00044 |
| ADCY10 | 16:35.6-35.9Mb | 0.117 | 0.00044 |
| PABPC4 | 1:76.7-78.7Mb | 0.117 | 0.00045 |
| DCLK1 | 12:9.7-10.6Mb | 0.117 | 0.00045 |
| TSHZ2 | 5:114.5-116.6Mb | 0.117 | 0.00045 |
| ARAP3 | 13:48.3-50Mb | 0.117 | 0.00045 |
| OSBP2 | 1:76.7-78.7Mb | 0.117 | 0.00046 |
| LRIG3 | 17:37-38.3Mb | 0.117 | 0.00046 |
| C7orf43 | 21:14.1-16.5Mb | 0.117 | 0.00046 |
| NR2C2 | 4:3.9-6.6Mb | 0.117 | 0.00046 |
| GRM7 | 12:57.9-62.4Mb | 0.117 | 0.00046 |
| DOCK9 | 19:7.1-7.9Mb | 0.117 | 0.00046 |
| PZP | 2:191-195.6Mb | 0.117 | 0.00046 |
| HCRTR2 | 7:116.2-117.6Mb | 0.117 | 0.00046 |
| MSI2 | 1:52.8-53.9Mb | 0.116 | 0.00047 |
| SFXN2 | 12:10.6-11.5Mb | 0.116 | 0.00047 |
| SMC2 | 22:22.6-22.8Mb | 0.116 | 0.00047 |
| RTTN | 2:201.1-201.9Mb | 0.116 | 0.00048 |
| CROCC | 1:76.7-78.7Mb | 0.116 | 0.00048 |
| RUNX1T1 | 7:63.7-64.4Mb | 0.116 | 0.00048 |
| BACH2 | 1:121.6-121.6Mb | 0.116 | 0.00049 |

| | | | |
|---|---|---|---|
| C2CD5 | 7:63.7-64.4Mb | 0.116 | 0.00049 |
| KCTD3 | 1:119.9-121.6Mb | 0.116 | 0.00049 |
| ADAM22 | 11:54.6-55.7Mb | 0.116 | 0.00049 |
| PAN2 | 1:206.1-207.3Mb | 0.116 | 0.00050 |
| KIF21B | 17:58.1-59.2Mb | 0.115 | 0.00051 |
| BCL9L | 11:55.7-56.2Mb | 0.115 | 0.00051 |
| EEF2K | 3:85.8-90.3Mb | 0.115 | 0.00051 |
| ICAM5 | 7:63.7-64.4Mb | 0.115 | 0.00052 |
| ACP4 | 1:117.9-119.9Mb | 0.115 | 0.00052 |
| ABCC3 | 1:40.7-42.6Mb | 0.115 | 0.00052 |
| ZNF479 | 13:48.3-50Mb | 0.115 | 0.00053 |
| ANO8 | 16:11.3-12.6Mb | 0.115 | 0.00053 |
| TP53 | 16:1.9-2.2Mb | 0.115 | 0.00053 |
| DOCK9 | 19:7.9-8.8Mb | 0.115 | 0.00053 |
| C7orf43 | 21:16.6-19.3Mb | 0.115 | 0.00053 |
| GRM8 | 12:55.4-56Mb | 0.115 | 0.00053 |
| ZNF7 | 19:27.6-29.5Mb | 0.115 | 0.00054 |
| RRAGB | 1:206.1-207.3Mb | 0.115 | 0.00054 |
| ICE1 | 22:22.8-23.4Mb | 0.115 | 0.00054 |
| NELL2 | 1:117.9-119.9Mb | 0.114 | 0.00055 |
| ESRP1 | 2:179.1-182.9Mb | 0.114 | 0.00055 |
| ASXL3 | 17:31.8-32.5Mb | 0.114 | 0.00056 |
| PARP10 | 11:54.6-55.7Mb | 0.114 | 0.00056 |
| SI | 11:99.1-102.8Mb | 0.114 | 0.00056 |
| MMP8 | 12:64.3-65.8Mb | 0.114 | 0.00056 |
| TGS1 | 17:31.8-32.5Mb | 0.114 | 0.00056 |
| DLEC1 | 1:119.9-121.6Mb | 0.114 | 0.00056 |
| DRD3 | 1:121.6-121.6Mb | 0.114 | 0.00057 |
| PTPDC1 | 12:10.6-11.5Mb | 0.114 | 0.00057 |
| ATP6V1A | 11:54.6-55.7Mb | 0.114 | 0.00057 |
| ADGRG1 | 1:117.9-119.9Mb | 0.114 | 0.00057 |
| STAG3 | 1:49.3-51.8Mb | 0.114 | 0.00057 |
| CAMSAP1 | 1:121.6-121.6Mb | 0.114 | 0.00057 |
| KDM1A | 22:22.2-22.6Mb | 0.114 | 0.00057 |
| FYCO1 | 4:87.3-88.7Mb | 0.114 | 0.00058 |
| POLR2A | 11:54.6-55.7Mb | 0.113 | 0.00058 |
| ANO8 | 16:9.6-11.3Mb | 0.113 | 0.00058 |
| ZNF516 | 17:19.2-19.7Mb | 0.113 | 0.00058 |
| FLG2 | 3:85.8-90.3Mb | 0.113 | 0.00058 |
| KDM1A | 22:21-21.6Mb | 0.113 | 0.00058 |
| UPF2 | 1:53.9-55.2Mb | 0.113 | 0.00059 |
| GRIK3 | 1:76.7-78.7Mb | 0.113 | 0.00059 |
| AMER3 | 6:105.1-107.8Mb | 0.113 | 0.00059 |
| DNM1P47 | 1:117.9-119.9Mb | 0.113 | 0.00059 |
| MUC3A | 1:121.6-121.6Mb | 0.113 | 0.00060 |
| TP53 | 16:1.4-1.9Mb | 0.113 | 0.00060 |
| F13A1 | 1:117.9-119.9Mb | 0.113 | 0.00060 |
| RB1 | 18:10.6-12.1Mb | 0.113 | 0.00060 |
| THBS1 | 22:21-21.6Mb | 0.113 | 0.00060 |
| RB1 | 18:1.5-3.5Mb | 0.113 | 0.00061 |
| PHF20L1 | 1:206.1-207.3Mb | 0.113 | 0.00061 |
| RB1 | 19:56.1-56.8Mb | 0.113 | 0.00061 |
| RRP12 | 1:39.7-40.7Mb | 0.113 | 0.00061 |
| TRIM71 | 12:10.6-11.5Mb | 0.113 | 0.00062 |
| PLEKHA4 | 1:76.7-78.7Mb | 0.113 | 0.00062 |
| FAM153A | 5:114.5-116.6Mb | 0.113 | 0.00062 |
| TP53 | 16:2.2-2.6Mb | 0.112 | 0.00062 |

| | | | |
|---|---|---|---|
| HCRTR2 | 5:100.7-107.2Mb | 0.112 | 0.00062 |
| NCOA6 | 11:55.7-56.2Mb | 0.112 | 0.00063 |
| ZNF224 | 12:10.6-11.5Mb | 0.112 | 0.00063 |
| GPRIN3 | 12:65.8-68.2Mb | 0.112 | 0.00063 |
| ABCC11 | 5:100.7-107.2Mb | 0.112 | 0.00064 |
| RRP12 | 1:206.1-207.3Mb | 0.112 | 0.00064 |
| C1QL4 | 1:121.6-121.6Mb | 0.112 | 0.00064 |
| T | 1:76.7-78.7Mb | 0.112 | 0.00064 |
| PWP2 | 1:121.6-121.6Mb | 0.112 | 0.00064 |
| PRUNE2 | 11:55.7-56.2Mb | 0.112 | 0.00065 |
| HEATR5A | 1:121.6-121.6Mb | 0.112 | 0.00065 |
| ATIC | 1:49.3-51.8Mb | 0.112 | 0.00065 |
| IFT172 | 7:63.7-64.4Mb | 0.112 | 0.00065 |
| POU6F2 | 6:6.7-10.1Mb | 0.112 | 0.00066 |
| ITGB4 | 1:170.1-171.8Mb | 0.112 | 0.00066 |
| SLC38A2 | 22:22.8-23.4Mb | 0.112 | 0.00066 |
| HUS1 | 12:10.6-11.5Mb | 0.112 | 0.00066 |
| LDLR | 1:119.9-121.6Mb | 0.112 | 0.00066 |
| ROCK2 | 2:92-92Mb | 0.112 | 0.00066 |
| CTAGE1 | 11:55.7-56.2Mb | 0.112 | 0.00067 |
| MMP2 | 17:39.1-40.1Mb | 0.112 | 0.00067 |
| COL2A1 | 13:48.3-50Mb | 0.112 | 0.00067 |
| ATIC | 1:53.9-55.2Mb | 0.112 | 0.00067 |
| SNX19 | 6:6.7-10.1Mb | 0.111 | 0.00067 |
| ARHGAP26 | 1:53.9-55.2Mb | 0.111 | 0.00067 |
| STAG3 | 1:52.8-53.9Mb | 0.111 | 0.00067 |
| DHX35 | 1:117.9-119.9Mb | 0.111 | 0.00067 |
| SPECC1 | 12:10.6-11.5Mb | 0.111 | 0.00067 |
| PLD5 | 17:38.4-39.1Mb | 0.111 | 0.00067 |
| KDM1A | 22:22.6-22.8Mb | 0.111 | 0.00068 |
| TANC2 | 17:39.1-40.1Mb | 0.111 | 0.00068 |
| HIST1H2BC | 11:55.7-56.2Mb | 0.111 | 0.00068 |
| THOC5 | 1:76.7-78.7Mb | 0.111 | 0.00069 |
| MYO18A | 12:9.7-10.6Mb | 0.111 | 0.00069 |
| BAZ1A | 13:48.3-50Mb | 0.111 | 0.00069 |
| MEGF8 | 19:27.6-29.5Mb | 0.111 | 0.00069 |
| XKR7 | 15:25.1-25.2Mb | 0.111 | 0.00069 |
| SHPRH | 7:18.8-22.4Mb | 0.111 | 0.00070 |
| VIL1 | 1:73.3-76.6Mb | 0.111 | 0.00070 |
| RIC1 | 16:28.8-29.5Mb | 0.111 | 0.00070 |
| ZCCHC6 | 1:121.6-121.6Mb | 0.111 | 0.00070 |
| RB1 | 18:3.5-6.4Mb | 0.111 | 0.00070 |
| HCRTR2 | 5:114.5-116.6Mb | 0.111 | 0.00071 |
| SLCO1C1 | 1:119.9-121.6Mb | 0.111 | 0.00071 |
| CREBRF | 5:114.5-116.6Mb | 0.111 | 0.00072 |
| RB1 | 16:19.1-20.6Mb | 0.110 | 0.00072 |
| PLEKHN1 | 12:55.4-56Mb | 0.110 | 0.00072 |
| ADAMTS10 | 12:57.9-62.4Mb | 0.110 | 0.00072 |
| DPH2 | 1:121.6-121.6Mb | 0.110 | 0.00072 |
| USP4 | 22:22.6-22.8Mb | 0.110 | 0.00072 |
| USP40 | 1:189.9-193.8Mb | 0.110 | 0.00073 |
| CD1E | 22:30.5-31.3Mb | 0.110 | 0.00073 |
| RB1 | 19:55.4-56.2Mb | 0.110 | 0.00073 |
| CASC1 | 2:179.1-182.9Mb | 0.110 | 0.00073 |
| METAP1D | 1:121.6-121.6Mb | 0.110 | 0.00074 |
| MRGPRF | 12:55.4-56Mb | 0.110 | 0.00074 |
| SCN3A | 15:71.5-72.9Mb | 0.110 | 0.00075 |

| | | | |
|---|---|---|---|
| WIPF1 | 1:76.7-78.7Mb | 0.110 | 0.00075 |
| KBTBD7 | 19:27.6-29.5Mb | 0.110 | 0.00075 |
| TYK2 | 12:56-56.4Mb | 0.110 | 0.00075 |
| CIART | 1:76.7-78.7Mb | 0.110 | 0.00076 |
| IWS1 | 7:56.6-57.4Mb | 0.110 | 0.00076 |
| ADTRP | 12:10.6-11.5Mb | 0.110 | 0.00076 |
| PADI1 | 5:116.6-120.3Mb | 0.110 | 0.00076 |
| VWCE | 13:50-51.9Mb | 0.110 | 0.00076 |
| ATIC | 1:51.7-52.8Mb | 0.110 | 0.00076 |
| DOCK6 | 2:127.9-130.2Mb | 0.110 | 0.00076 |
| STAG3 | 1:51.7-52.8Mb | 0.110 | 0.00076 |
| CNOT10 | 12:9.7-10.6Mb | 0.110 | 0.00076 |
| CASK | 13:48.3-50Mb | 0.110 | 0.00076 |
| ADGRG1 | 1:121.6-121.6Mb | 0.110 | 0.00076 |
| MSI2 | 1:40.7-42.6Mb | 0.110 | 0.00077 |
| PLCL2 | 6:105.1-107.8Mb | 0.110 | 0.00077 |
| ZNF599 | 12:62.5-64.2Mb | 0.110 | 0.00077 |
| TP53 | 16:3.1-3.5Mb | 0.109 | 0.00077 |
| MPRIP | 1:121.6-121.6Mb | 0.109 | 0.00077 |
| ITPRIPL1 | 11:54.6-55.7Mb | 0.109 | 0.00077 |
| ADGRL1 | 7:63.7-64.4Mb | 0.109 | 0.00077 |
| ZC3H13 | 1:76.7-78.7Mb | 0.109 | 0.00078 |
| ARHGAP33 | 1:119.9-121.6Mb | 0.109 | 0.00078 |
| T | 1:117.9-119.9Mb | 0.109 | 0.00078 |
| ARMC5 | 19:29.5-32.5Mb | 0.109 | 0.00078 |
| RABGGTA | 13:48.3-50Mb | 0.109 | 0.00078 |
| RB1 | 16:0.9-1.4Mb | 0.109 | 0.00078 |
| TP53 | 16:2.6-3.1Mb | 0.109 | 0.00079 |
| SOX5 | 12:65.8-68.2Mb | 0.109 | 0.00079 |
| RB1 | 9:21.5-25.8Mb | 0.109 | 0.00079 |
| CNTNAP3 | 1:121.6-121.6Mb | 0.109 | 0.00079 |
| ADAMTS13 | 1:121.6-121.6Mb | 0.109 | 0.00079 |
| WDR36 | 5:114.5-116.6Mb | 0.109 | 0.00079 |
| WDFY3 | 17:31.8-32.5Mb | 0.109 | 0.00079 |
| SPATA31D1 | 1:76.7-78.7Mb | 0.109 | 0.00080 |
| CNTNAP2 | 12:9.7-10.6Mb | 0.109 | 0.00080 |
| SIGLEC12 | 12:10.6-11.5Mb | 0.109 | 0.00080 |
| ZNF615 | 1:121.6-121.6Mb | 0.109 | 0.00080 |
| TRPM2 | 1:53.9-55.2Mb | 0.109 | 0.00080 |
| NUDT22 | 2:127.9-130.2Mb | 0.109 | 0.00080 |
| LTBP1 | 1:39.7-40.7Mb | 0.109 | 0.00080 |
| TPGS2 | 1:119.9-121.6Mb | 0.109 | 0.00080 |
| PWP2 | 1:119.9-121.6Mb | 0.109 | 0.00081 |
| KIDINS220 | 12:54-54.3Mb | 0.109 | 0.00081 |
| BUB1 | 1:206.1-207.3Mb | 0.109 | 0.00081 |
| MTMR11 | 11:54.6-55.7Mb | 0.109 | 0.00081 |
| KDM1A | 22:21.6-22.2Mb | 0.109 | 0.00082 |
| TNK2 | 1:62.2-63.8Mb | 0.108 | 0.00083 |
| BPTF | 7:53.5-55.7Mb | 0.108 | 0.00083 |
| AKT1 | 11:99.1-102.8Mb | 0.108 | 0.00083 |
| IGSF9B | 22:22.8-23.4Mb | 0.108 | 0.00083 |
| HERC2P3 | 1:76.7-78.7Mb | 0.108 | 0.00083 |
| HHIPL2 | 12:10.6-11.5Mb | 0.108 | 0.00084 |
| LY9 | 3:49.9-50.6Mb | 0.108 | 0.00084 |
| GTF2A1 | 13:48.3-50Mb | 0.108 | 0.00084 |
| COL27A1 | 1:117.9-119.9Mb | 0.108 | 0.00085 |
| RB1 | 19:48.9-49.3Mb | 0.108 | 0.00085 |

| | | | |
|---|---|---|---|
| PCDHA1 | 6:105.1-107.8Mb | 0.108 | 0.00085 |
| SLC8A3 | 11:77.1-78.6Mb | 0.108 | 0.00085 |
| ZNF615 | 1:76.7-78.7Mb | 0.108 | 0.00085 |
| CCDC18 | 1:121.6-121.6Mb | 0.108 | 0.00086 |
| CEP250 | 7:63.7-64.4Mb | 0.108 | 0.00087 |
| MMP2 | 17:48.3-48.9Mb | 0.108 | 0.00087 |
| CYTH3 | 17:15.4-16.1Mb | 0.108 | 0.00087 |
| SLC24A1 | 12:55.4-56Mb | 0.108 | 0.00087 |
| MIB1 | 13:48.3-50Mb | 0.108 | 0.00087 |
| ZNF534 | 11:99.1-102.8Mb | 0.108 | 0.00087 |
| HECTD4 | 1:55.2-58.6Mb | 0.108 | 0.00087 |
| TSEN2 | 1:121.6-121.6Mb | 0.108 | 0.00088 |
| FAM171B | 12:55.4-56Mb | 0.108 | 0.00088 |
| PMM1 | 4:3.9-6.6Mb | 0.108 | 0.00088 |
| FAM171A1 | 6:11.5-14.6Mb | 0.108 | 0.00088 |
| DSCAML1 | 19:7.1-7.9Mb | 0.108 | 0.00088 |
| NLN | 2:191-195.6Mb | 0.108 | 0.00088 |
| ZMIZ1 | 19:27.6-29.5Mb | 0.108 | 0.00088 |
| KIFAP3 | 4:143.6-146.1Mb | 0.108 | 0.00089 |
| ATP2B1 | 1:206.1-207.3Mb | 0.108 | 0.00089 |
| ANKRD35 | 1:121.6-121.6Mb | 0.108 | 0.00089 |
| TIAM1 | 1:121.6-121.6Mb | 0.108 | 0.00089 |
| TTC7A | 22:22.6-22.8Mb | 0.108 | 0.00089 |
| HNRNPDL | 1:121.6-121.6Mb | 0.107 | 0.00089 |
| MSI2 | 1:55.2-58.6Mb | 0.107 | 0.00089 |
| PTPN22 | 7:112.3-116.3Mb | 0.107 | 0.00089 |
| ABHD16A | 11:105.2-108.6Mb | 0.107 | 0.00089 |
| RSC1A1 | 5:114.5-116.6Mb | 0.107 | 0.00090 |
| PSMD1 | 1:76.7-78.7Mb | 0.107 | 0.00090 |
| TP53 | 16:0-0.5Mb | 0.107 | 0.00090 |
| UBE4A | 12:10.6-11.5Mb | 0.107 | 0.00090 |
| PAN2 | 12:27-29Mb | 0.107 | 0.00090 |
| PAPD7 | 2:179.1-182.9Mb | 0.107 | 0.00090 |
| CCDC191 | 1:53.9-55.2Mb | 0.107 | 0.00091 |
| KDM3B | 7:112.3-116.3Mb | 0.107 | 0.00091 |
| CEMIP | 6:105.1-107.8Mb | 0.107 | 0.00091 |
| OR5H15 | 7:38.7-42.8Mb | 0.107 | 0.00091 |
| MMP8 | 12:65.8-68.2Mb | 0.107 | 0.00091 |
| FOXP1 | 12:10.6-11.5Mb | 0.107 | 0.00091 |
| ZNF394 | 1:117.9-119.9Mb | 0.107 | 0.00092 |
| IGSF9B | 22:21-21.6Mb | 0.107 | 0.00092 |
| ARID1B | 1:47-49.2Mb | 0.107 | 0.00092 |
| ALMS1 | 4:184.3-185.6Mb | 0.107 | 0.00092 |
| ABCC12 | 1:76.7-78.7Mb | 0.107 | 0.00093 |
| DOT1L | 4:3.9-6.6Mb | 0.107 | 0.00093 |
| ZFHX3 | 11:77.1-78.6Mb | 0.107 | 0.00093 |
| SYTL3 | 5:114.5-116.6Mb | 0.107 | 0.00093 |
| JARID2 | 21:39.4-41.9Mb | 0.107 | 0.00093 |
| LRRC37A3 | 12:65.8-68.2Mb | 0.107 | 0.00093 |
| MIER1 | 1:119.9-121.6Mb | 0.107 | 0.00093 |
| SGIP1 | 12:65.8-68.2Mb | 0.107 | 0.00093 |
| ABHD3 | 7:116.2-117.6Mb | 0.107 | 0.00094 |
| RB1 | 19:51-51.4Mb | 0.107 | 0.00094 |
| KIF21A | 11:60.4-61.4Mb | 0.107 | 0.00094 |
| PAPSS2 | 12:55.4-56Mb | 0.107 | 0.00094 |
| STIL | 1:40.7-42.6Mb | 0.107 | 0.00094 |
| LETMD1 | 1:76.7-78.7Mb | 0.107 | 0.00095 |

| | | | |
|---|---|---|---|
| MFHAS1 | 22:22.8-23.4Mb | 0.107 | 0.00095 |
| MRGPRX1 | 17:37-38.3Mb | 0.107 | 0.00095 |
| PAX3 | 1:76.7-78.7Mb | 0.107 | 0.00095 |
| CROCC | 1:73.3-76.6Mb | 0.107 | 0.00095 |
| PHF2 | 21:14.1-16.5Mb | 0.107 | 0.00095 |
| ABCC12 | 1:73.3-76.6Mb | 0.107 | 0.00095 |
| PTPRG | 17:60.1-61.6Mb | 0.107 | 0.00095 |
| MCM3AP | 6:6.7-10.1Mb | 0.107 | 0.00095 |
| FBLN1 | 1:117.9-119.9Mb | 0.107 | 0.00095 |
| ZNF608 | 2:201.1-201.9Mb | 0.107 | 0.00095 |
| FAM47B | 21:16.6-19.3Mb | 0.107 | 0.00095 |
| COL5A2 | 17:19.2-19.7Mb | 0.107 | 0.00096 |
| BRD9 | 19:27.6-29.5Mb | 0.106 | 0.00096 |
| MYO15A | 17:36-36.9Mb | 0.106 | 0.00096 |
| CFLAR | 1:121.6-121.6Mb | 0.106 | 0.00096 |
| CFAP43 | 12:56-56.4Mb | 0.106 | 0.00096 |
| FKBP5 | 7:63.7-64.4Mb | 0.106 | 0.00096 |
| KCNIP1 | 4:3.9-6.6Mb | 0.106 | 0.00097 |
| ADAM22 | 11:55.7-56.2Mb | 0.106 | 0.00097 |
| LTBP1 | 1:206.1-207.3Mb | 0.106 | 0.00097 |
| GRM8 | 12:54-54.3Mb | 0.106 | 0.00098 |
| KIAA1551 | 7:65.5-66.6Mb | 0.106 | 0.00098 |
| HSPA14 | 17:37-38.3Mb | 0.106 | 0.00098 |
| FAM98C | 17:31.8-32.5Mb | 0.106 | 0.00098 |
| CAND2 | 17:58.1-59.2Mb | 0.106 | 0.00098 |
| ECEL1 | 5:100.7-107.2Mb | 0.106 | 0.00099 |
| MUC3A | 1:119.9-121.6Mb | 0.106 | 0.00099 |
| SPICE1 | 1:40.7-42.6Mb | 0.106 | 0.00099 |
| PADI2 | 1:73.3-76.6Mb | 0.106 | 0.00099 |
| ANKRD30A | 4:87.3-88.7Mb | 0.106 | 0.00100 |
| DNM1P47 | 1:119.9-121.6Mb | 0.106 | 0.00100 |
| TPCN1 | 1:117.9-119.9Mb | 0.106 | 0.00100 |
| CNOT10 | 12:10.6-11.5Mb | 0.106 | 0.00100 |
| ETAA1 | 2:179.1-182.9Mb | 0.106 | 0.00100 |

### 3 B-spline functions

Marginal and joint probabilities are estimated before MI or CMI calculation. For a variable $X$ with observed values $x_1, \ldots, x_N$, these values are normalized between 0 and 1:

$$\hat{x} = \frac{x - x_{\min}}{x_{\max} - x_{\min}}$$

The range between 0 and 1 is evenly divided into $B$ bins of equal size, represented by $\mathcal{X}_1, ..., \mathcal{X}_B$. B-spline functions [1] are used to estimate the probability mass function $p(\mathcal{X})$. A knot vector $t_i$ (Fig. 1) is defined based upon the bin number $i$ and spline order $s$ where $i, s \in \mathbb{Z}$ and $1 \leq i, s \leq B$:

$$t_i := \begin{cases} 0, & \text{if } i < s \\ \frac{i-s+1}{B-s+1}, & \text{if } s \leq i \leq B-1 \\ 1, & \text{if } i > B-1 \end{cases}$$

Intuitively, the knot vector splits the range from 0 to 1 into $B - s + 1$ evenly spaced segments (Fig. 1).

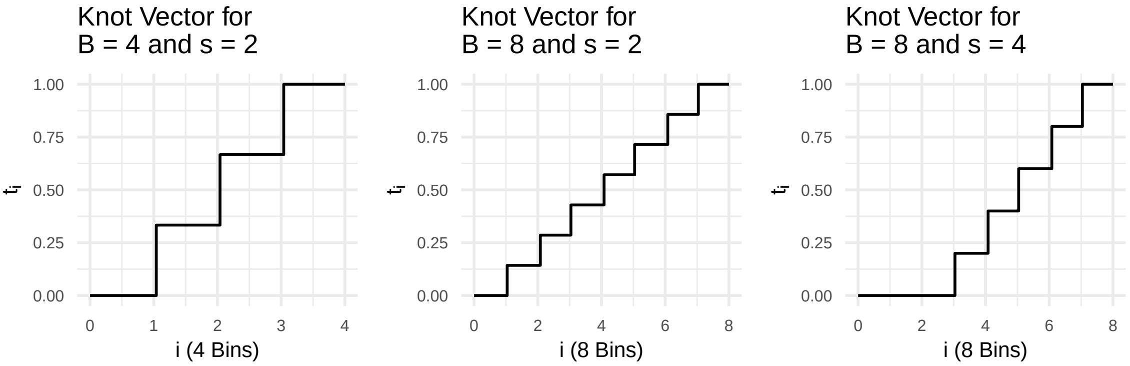


Figure 1: Illustration of knot vectors with varying choices for bin number and spline order.

B-spline basis functions $B_{i,s}(\hat{x})$ are constructed in order to map $\hat{x}$ data values into appropriate bins. The B-spline functions are recursively constructed as follows:

$$B_{i,1}(\hat{x}) = \begin{cases} 1, & \text{if } t_i \leq \hat{x} < t_{i+1} \\ 0, & \text{otherwise} \end{cases}$$

$$B_{i,s}(\hat{x}) = \frac{\hat{x} - t_i}{t_{i+s-1} - t_i} B_{i,s-1}(\hat{x}) + \frac{t_{i+s} - \hat{x}}{t_{i+s} - t_{i+1}} B_{i+1,s-1}(\hat{x})$$

B-spline basis functions with $s > 1$ can assign each data point $\hat{x}$ into multiple bins (Fig. 2), thus providing an estimation of association that is more robust to impact from outliers [1]. The probability mass function $p(\mathcal{X})$ is then estimated as:

$$p(\mathcal{X}_i) = \frac{1}{N} \sum_{v=1}^{N} B_{i,s}(\hat{x}_v)$$

where $B_{i,s}(\hat{x}_v)$ represents the probability of $\hat{x}_v$ belonging to bin $i$ given spline order $s$. The sum of such B-spline functions over all $N$ observed data points is used to estimate the probability of bin $i$, $p(\mathcal{X}_i)$.

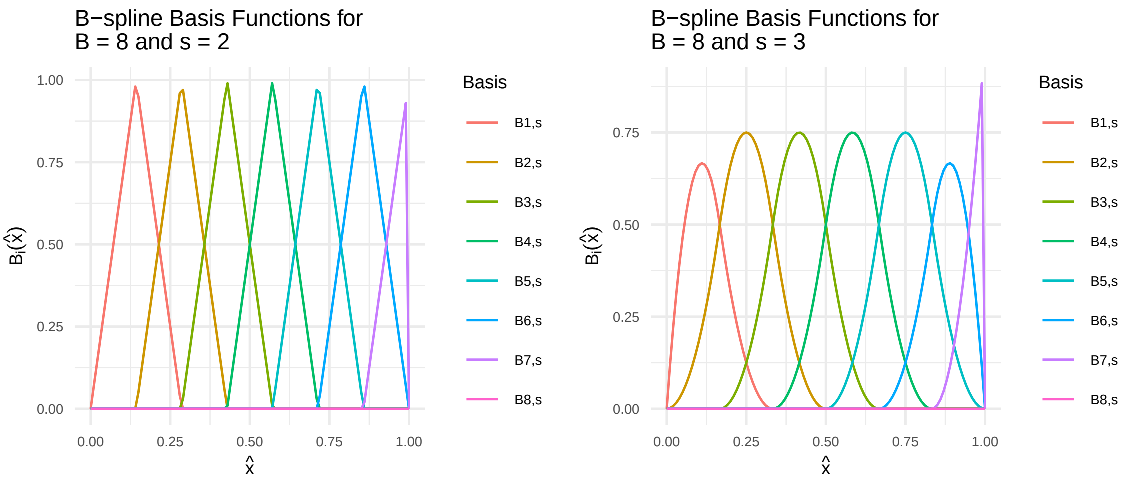


Figure 2: Illustration of B-spline basis functions with different spline order.

### 4 Impact of bin number and spline order on association strength and statistical significance

We used TCGA BLCA (Urothelial Bladder Carcinoma) data from Section 3.1 of the main manuscript to demonstrate the impact of different hyperparameter settings on MI/CMI association strength and statistical significance. We randomly selected 10,000 mutation-SCNA pairs from all possible combinations (Fig. 3). The MI between mutation and SCNA, CMI between mutation and SCNA conditioned on tumor purity (termed $\text{CMI}_p$), and CMI between mutation and SCNA conditioned on tumor purity and mutation burden (termed $\text{CMI}_{p,m}$) were estimated (Fig. 3). We observed a strong positive correlation amongst the association values obtained using different bins and spline orders (Fig. 3). We also observed that an increased spline order tended to cause greater variance in the magnitude of association strengths (Fig. 3).

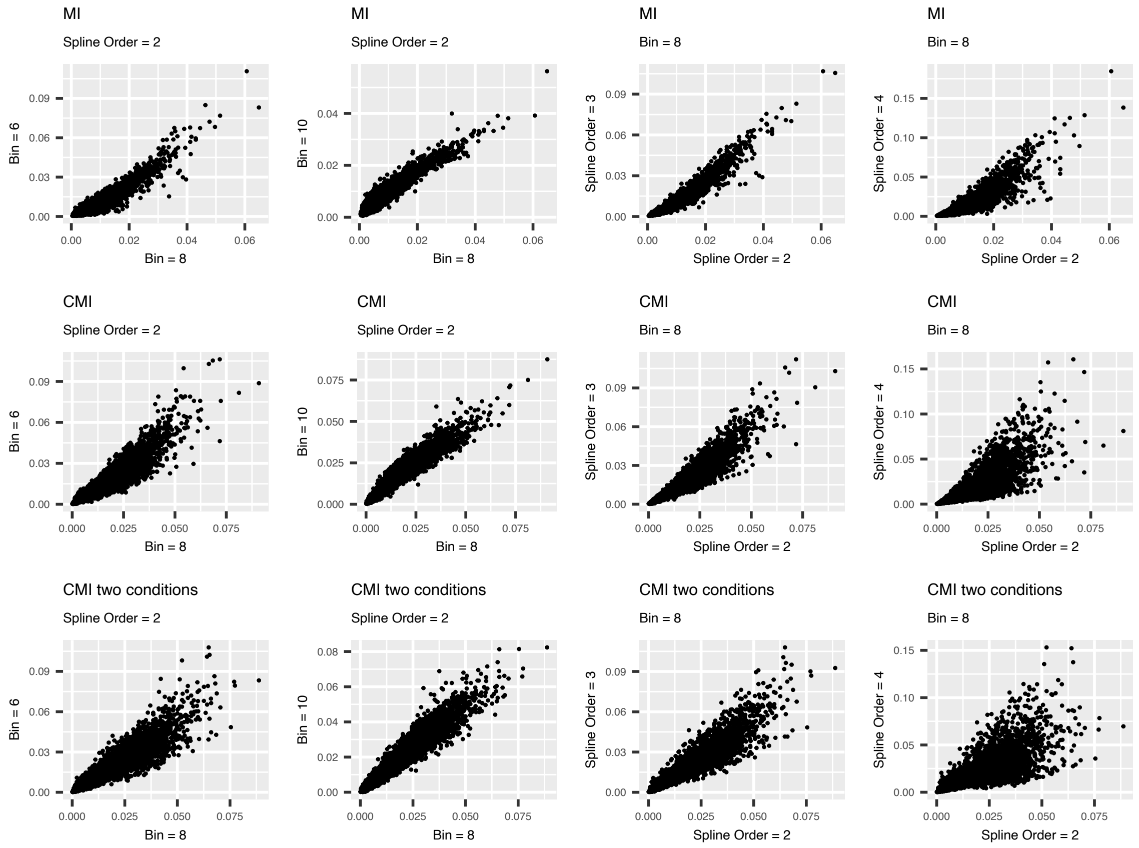


Figure 3: Impact of bin number and spline order on association strength for randomly selected mutation-SCNA pairs. 10,000 mutation-SCNA pairs were randomly selected. MI, $\text{CMI}_p$, and $\text{CMI}_{p,m}$ were evaluated between these 10,000 randomly selected pairs.

The statistical significance of MI, $\text{CMI}_p$, and $\text{CMI}_{p,m}$ was also estimated. We selected the top 500 mutation-SCNA pairs with the highest values for these metrics, using a bin size of 8 and a spline order of 2 (Fig. 4). We further evaluated the associations and statistical significance of these pairs by changing the bin sizes and spline orders to determine whether these changes would impact the association strength and statistical significance. Our findings revealed that increasing the number of bins generally reduces the magnitude of association values (for MI, $\text{CMI}_p$, and $\text{CMI}_{p,m}$, Fig. 4). In contrast, increasing the spline order only increases the association values for MI (but not for $\text{CMI}_p$, $\text{CMI}_{p,m}$, Fig. 4). Importantly, the selection of spline order should match the bin size, since an increase in spline order may increase the association values obtained from bootstrapping and increase the corresponding p value as well. We suggest that the spline order should be smaller than half of the bin size. In general, altering the bin sizes and spline orders did not significantly affect the statistical significance of MI, $\text{CMI}_p$, and $\text{CMI}_{p,m}$. These observations are consistent with previous research suggesting that B-spline functions are robust methods for MI estimation [1]. Here we demonstrated that the same B-spline functions can also be applied for

CMI estimation.

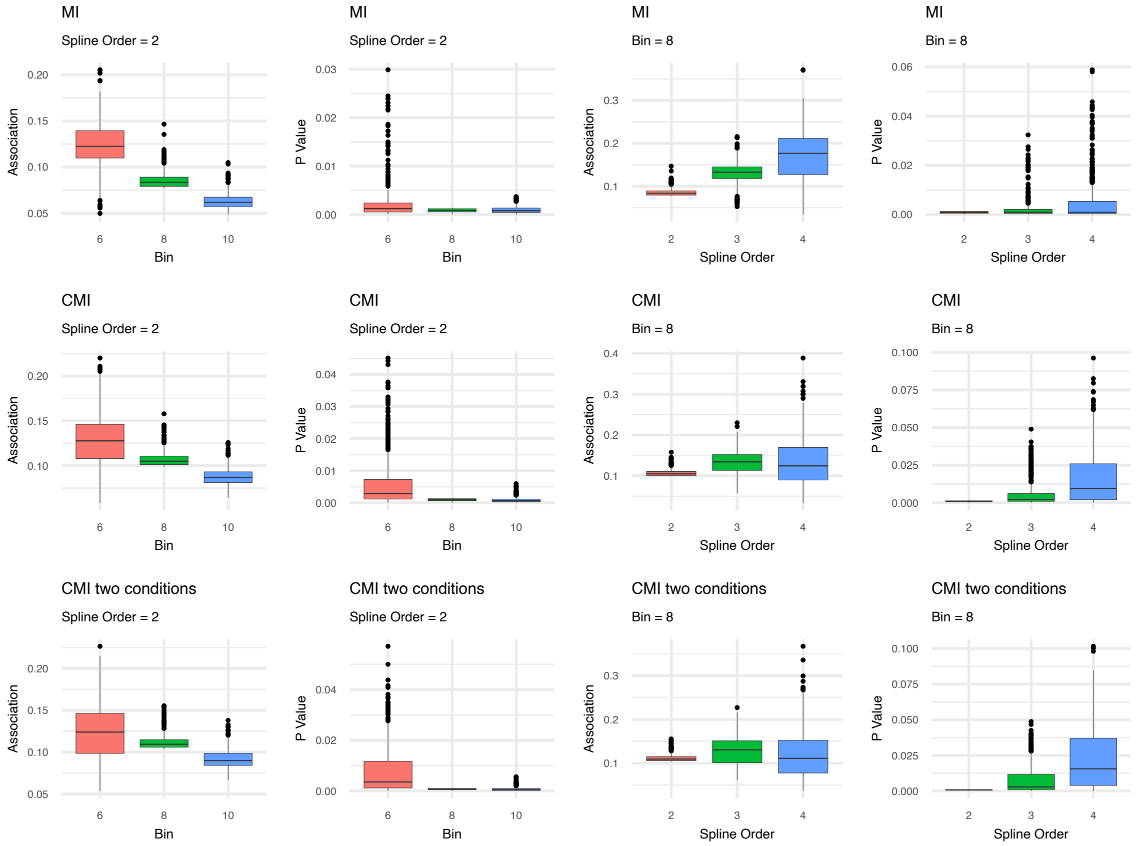


Figure 4: Impact of bin number and spline order on association strength and statistical significance in the top 500 mutation-SCNA pairs. Top 500 mutation-SCNA pairs were selected for MI, $\mathrm{CMI}_p$ and $\mathrm{CMI}_{p,m}$ obtained from bin of 8 and spline order of 2. Then MI, $\mathrm{CMI}_p$ and $\mathrm{CMI}_{p,m}$ of these mutation-SCNA pairs were estimated again using different bins and spline orders

### 5 Comparison between Correlations/Linear Regression and MI/CMI

Pearson/Spearman correlations and linear regressions are commonly used methods to measure associations in biological data. Using the lung cancer tumor microenvironment cell fraction dataset from Section 3.2, Pearson and Spearman correlations along with their associated p values were calculated for every pair of cell types.

MI/CMI was calculated for every pair of cell types using hyperparameters bin = 8 and spline order = 2. CMI was calculated by conditioning on tumor purity, or the malignant cell fraction within the tumor microenvironment. MI/CMI values were lastly signed by multiplying with the sign of the Spearman correlation between each corresponding pair of cell fraction vectors, in order to provide information on whether the cell types tend to associate together or dissociate from one another in the tumor microenvironment.

Linear regression was performed (either with or without tumor purity as an additional covariate) as follows: for every pair of cell types $i, j$ with corresponding cell fraction vectors $F_i, F_j$ (where the length of $F_i, F_j$ equals the number of samples), a linear regression was performed as either

$$F_i = m_1 * F_j + b$$

or

$$F_i = m_1 * F_j + m_2 * P + b$$

where $P$ is the vector of tumor purities (malignant cell fractions). Resulting slopes $m_1$ were scaled by multiplying with $\sigma_j/\sigma_i$, where $\sigma_i, \sigma_j$ are the sample standard deviations for $F_i, F_j$ respectively. This scaling procedure adjusts the magnitude of the $m_1$ slope terms to range between $-1$ and $+1$, and produces a result equivalent to the Pearson correlation in the case where tumor purity is not included as a covariate in the linear regression. The p values associated with each $m_1$ slope term were recorded as well.

Finally, p values from each method were adjusted according to the Benjamini-Hochberg protocol, and an FDR threshold of 0.1 was set to filter out statistically insignificant results. Shown in the figure below is a side-by-side illustration of the statistically significant cell pair associations discovered by each of the six methods:

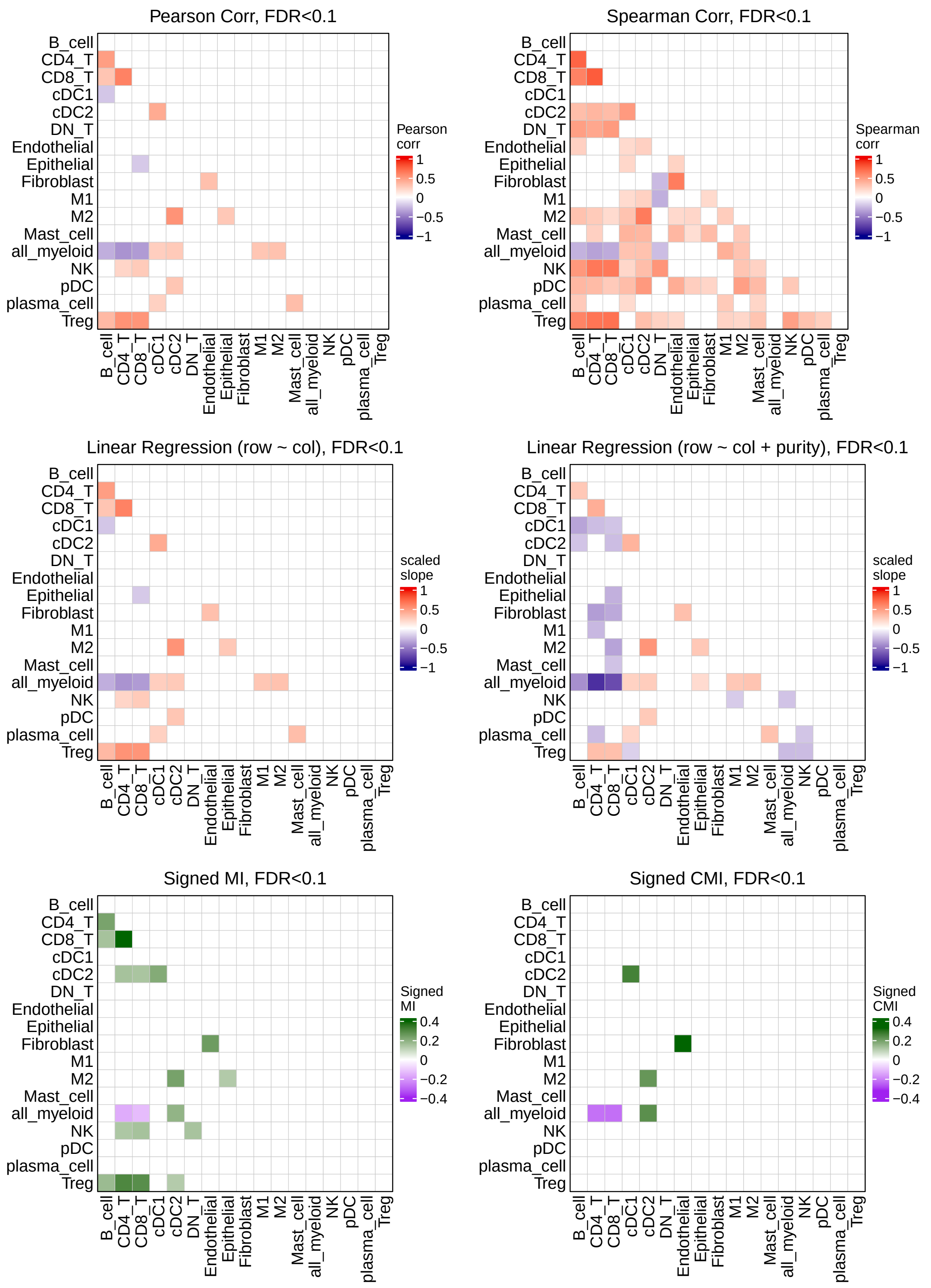


Figure 5: Statistically significant cell pair associations in lung cancer tumor microenvironment, as calculated by Pearson correlation (upper left), Spearman correlation (upper right), linear regression without tumor purity covariate (middle left), linear regression with tumor purity covariate (middle right), signed MI (lower left), signed CMI (lower right)

Spearman correlation discovered more statistically significant cell pair associations than Pearson correlation. Furthermore, the Pearson correlation heatmap is identical to the linear regression heatmap (without inclusion of tumor purity as a covariate). The set of significant cell pair associations discovered by Pearson correlation is highly similar, but not identical, to the set of significant cell pair associations discovered by MI. The significant cell pair associations discovered by MI are a strict subset of those discovered by Spearman correlation, and the significant cell pair associations discovered by CMI are a strict subset of those discovered by MI. That there are fewer statistically significant cell pair associations discovered by CMI compared to MI suggests that some of the cell pair associations discovered by MI may be driven (at least in part) by tumor purity, an external confounding variable.

On the other hand, there were a few cell pairs whose magnitude of signed CMI (conditioned on tumor purity) was *larger* than the corresponding magnitude of signed MI, suggesting that the cell pair association in these scenarios may have been obscured by the tumor purity covariate. Notable examples included: endothelial vs fibroblast (see Figure 6 first row), all myeloid cells vs CD4$^+$ T cells (see Figure 6 second row), and all myeloid cells vs CD8$^+$ T cells (see Figure 6 third row):

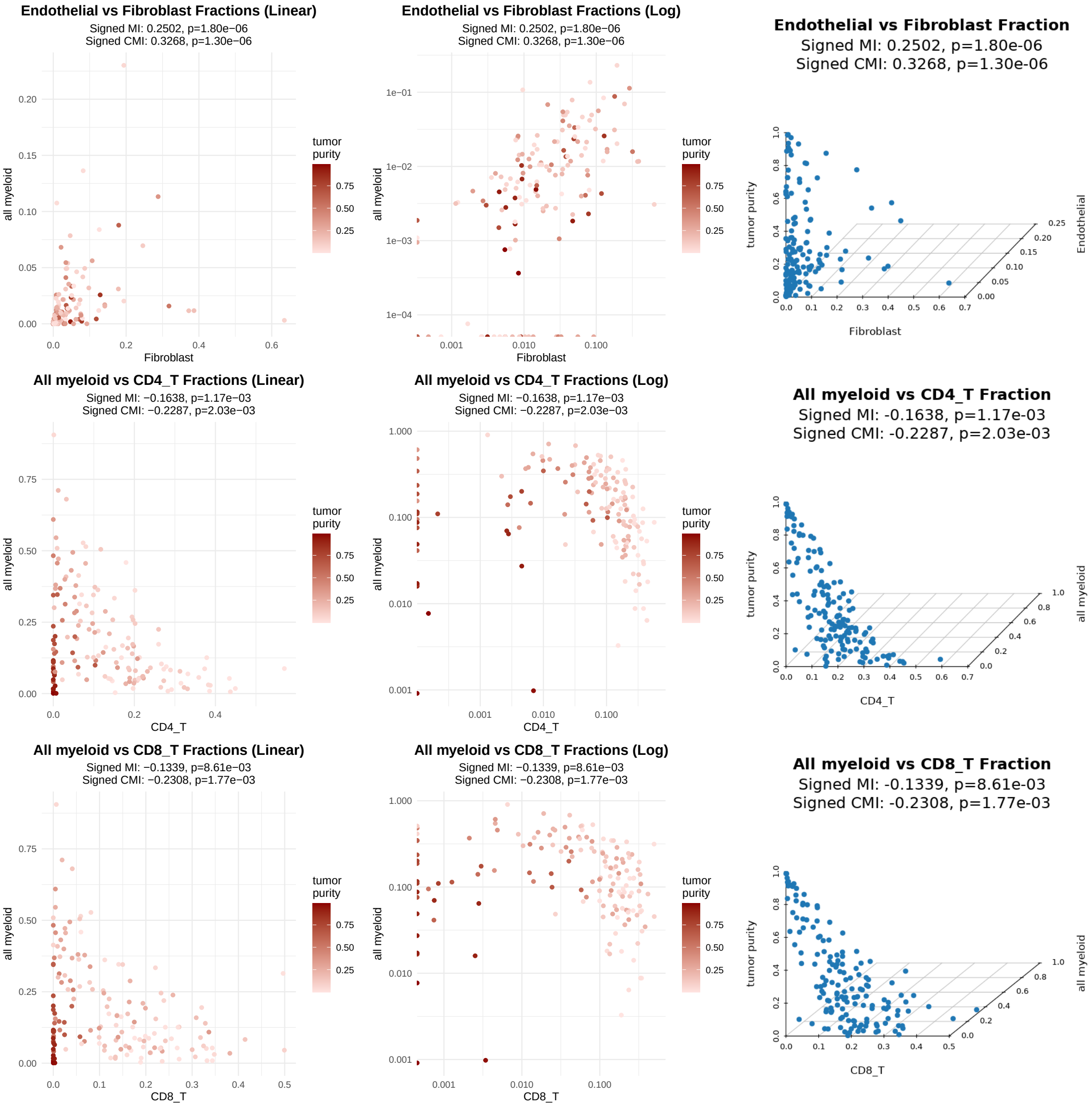


Figure 6: Three cell pairs with CMI > MI. First column: plotted with linear scale axes, second column: plotted with log scale axes, third column: plotted in three dimensions with tumor purity on the $z$ axis. MI in two dimensions is likely weakened by a multitude of high-purity (dark red) data points gathered near the origin, but CMI with three dimensions of information is likely better able to detect a strong magnitude of positive (or negative) association at varying levels of tumor purity.

We also observed more statistically significant cell pair associations discovered with Spearman correlation as opposed to MI. For this comparison, it is essential to recall that Spearman correlation and MI are fundamentally different measurements. For instance, consider two vectors $X, Y$ defined as

$$X = \{1, 2, 3, 4, 5, 6, 7, 8\}$$
$$Y = \{0, 0, 0, 0, 1, 1, 1, 1\}$$

The Spearman correlation between $X$ and $Y$ is 1, as there is a perfect monotonically increasing relationship between the two vectors. However, if $X$ and $Y$ are considered to be discrete random variables with probability mass function uniformly distributed over each of the eight values, then $I(X;Y)$ normalized with respect to $H(X)$ is only $\frac{1}{3}$. In other words, the MI between $X$ and $Y$ is relatively low because the value of $Y$ does not reveal much information about the value of $X$.

An example with ground-truth illustrating a similar trend can be found in the classic Waffle Houses dataset [7], which contains data on the number of Waffle House restaurants, median age of marriage, and divorce rates for every state within the United States. The basic premise of this problem is that there appears to be a statistically significant correlation between per-state number of Waffle House restaurants and divorce rate (see Figure 7). However, this correlation is confounded by an external variable: the median age of marriage in each state. Most Waffle House restaurants are located in the southern portion of the United States, which on average have lower median ages of marriage and higher divorce rates.

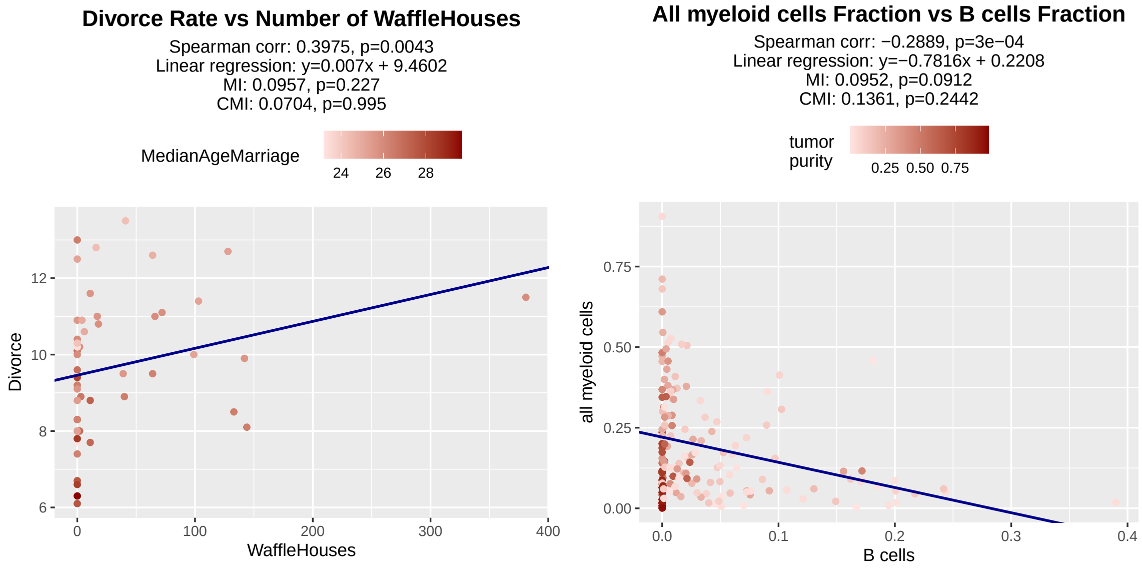


Figure 7: (Left) Per-state divorce rate vs number of Waffle House restaurants, colored by median age of marriage (Right) All myeloid cell fraction vs B cell fraction, colored by tumor purity

While the Spearman correlation for this data is moderately large and reaches statistical significance, the MI has a p value of 0.227. The absence of statistical significance for MI is likely driven by the abundance of data points lined up along the y-axis, representing states that have zero Waffle House locations but non-zero divorce rates. In the same manner as the toy example above, the abundance of these data points reduces the amount of information which the x-axis variable reveals about the y-axis variable, thereby resulting in a smaller MI that fails to reach statistical significance.

Furthermore, the CMI between divorce rates and number of Waffle House restaurants (conditioned on median age of marriage) has an even smaller magnitude compared to MI and an even larger p value. This is consistent with previous expectations – that median age of marriage is a confounding variable in the association between divorce rate and number of Waffle House restaurants. Overall, this example illustrates how `conMItion` is capable of filtering out spurious correlations driven by external confounding variables.

Returning to the lung cancer tumor microenvironment cell fraction dataset, a similar pattern may help to explain why there are far more statistically significant cell pair associations discovered by Spearman correlation than MI. Similar to the Waffle Houses dataset, this dataset has an abundance of 0 entries, representing tumor biopsy samples that had 0 counts of a particular cell type. (This often arises in high-purity tumor samples that are predominantly composed of malignant cells). See the right panel of Figure 7, illustrating the relationship between all myeloid cell fraction vs B cell fraction. Similar to the Waffle House example, there are many data points lined up against the y-axis, contributing to a Spearman correlation that is statistically significant but MI with larger, statistically insignificant p values. Furthermore, tumor purity is a potential confounding variable in this scenario, since the high-purity (dark red) data points are associated with having lower B cell fractions. This likely explains why the CMI for this cell pair relationship has an even larger p value, similar to what was observed with the Waffle House dataset.

To quantify this pattern more formally and across all cell type pairs being examined, consider an arbitrary pair of cell types, cell type $i$ and cell type $j$, with corresponding cell fraction vectors $F_i$ and $F_j$. $F_i$ and $F_j$ have length equal to the number of samples. For every sample $s$, we can define its sample tuple $(F_{is}, F_{js})$ to represent the fraction of cell type $i$ and $j$ in sample $s$. For each cell type pair $i, j$, we define the number of its *unique intercepts* as the number of unique sample tuples satisfying the form $(0, F_{js})$ where $F_{js} \neq 0$ or $(F_{is}, 0)$ where $F_{is} \neq 0$. We hypothesized that cell type pairs which fail to achieve statistically significant MI are more likely to have a greater number of unique intercepts. This can be visualized in the scatterplots and violin plot below:

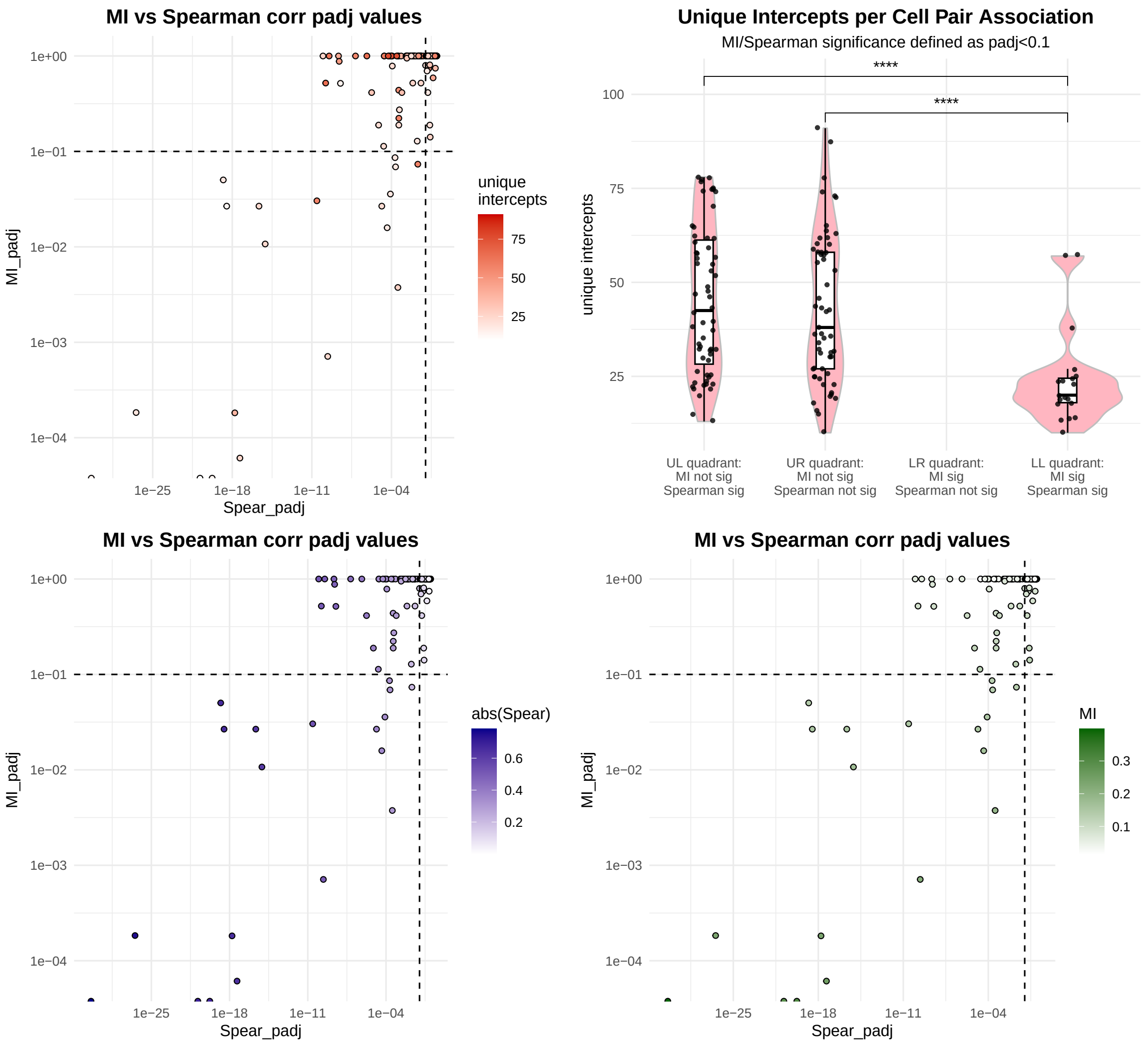


Figure 8: MI vs Spearman correlation adjusted p values, colored by number of unique intercepts (upper left) or absolute value of Spearman correlation (lower left) or MI (lower right); Distributions of number of unique intercepts per cell pair association, grouped by quadrant (upper right)

Each point in the three scatterplots above represents a cell pair, with its x-axis value representing the Spearman correlation adjusted p value and its y-axis value representing the MI adjusted p value. Each cell pair data point is furthermore colored by its quantity of unique intercepts (upper left), absolute value of Spearman correlation (lower left), or MI (lower right). Dashed horizontal and dashed vertical lines are located at $x = 0.1$ and $y = 0.1$ to indicate the FDR thresholds. As expected, the data points further to the left of the scatterplot (smaller Spearman correlation adjusted p value) have a larger magnitude of Spearman correlation (see lower left scatterplot), and the data points further to the bottom of the scatterplot (smaller MI adjusted p value) have a larger magnitude of MI (see lower right scatterplot). The location of points can be interpreted as follows:

- Upper left (UL) quadrant: cell pair associations that are significant according to Spearman but not according to MI
- Upper right (UR) quadrant: cell pair associations that are not significant according to both methods
- Lower right (LR) quadrant: cell pair associations that are significant according to MI but not according to Spearman
- Lower left (LL) quadrant: cell pair associations that are significant according to both methods

Interestingly, there are no data points in the lower right quadrant, indicating that the statistically significant cell pair associations identified by MI are a strict subset of those identified by Spearman. Most of the dark red points (cell pairs with high numbers of unique intercepts) are located in the upper left and upper right quadrants, but this pattern can be examined more thoroughly in the violin plot on the upper right. The cell pair associations that are classified as significant by both MI and Spearman (LL quadrant) have significantly fewer unique intercepts than cell pairs in the upper left quadrant (significant according to Spearman but not MI) and cell pairs in the upper right quadrant (not significant according to both methods). Overall, this result is highly consistent with the intuition built from the previous three examples – that an abundance of data points where one random variable is held constant while the other random variable varies reduces the magnitude and statistical significance of MI without having the same impact on Spearman correlation.
